# Global and Local Mutations in Bangladeshi SARS-CoV-2 Genomes

**DOI:** 10.1101/2020.08.25.267658

**Authors:** Md. Mahbub Hasan, Rasel Das, Md. Rasheduzzaman, Md Hamed Hussain, Nazmul Hasan Muzahid, Asma Salauddin, Meheadi Hasan Rumi, S M Mahbubur Rashid, AMAM Zonaed Siddiki, Adnan Mannan

## Abstract

Corona Virus Disease-2019 (COVID-19) warrants comprehensive investigations of publicly available Severe Acute Respiratory Syndrome-CoronaVirus-2 (SARS-CoV-2) genomes to gain new insight about their epidemiology, mutations and pathogenesis. Nearly 0.4 million mutations were identified so far in ∼60,000 SARS-CoV-2 genomic sequences. In this study, we compared 207 of SARS-CoV-2 genomes reported from different parts of Bangladesh and their comparison with 467 globally reported sequences to understand the origin of viruses, possible patterns of mutations, availability of unique mutations, and their apparent impact on pathogenicity of the virus in victims of Bangladeshi population. Phylogenetic analyses indicates that in Bangladesh, SARS-CoV-2 viruses might arrived through infected travelers from European countries, and the GR clade was found as predominant in this region. We found 2602 mutations including 1602 missense mutations, 612 synonymous mutations, 36 insertions and deletions with 352 other mutations types. In line with the global trend, D614G mutation in spike glycoprotein was predominantly high (95.6%) in Bangladeshi isolates. Interestingly, we found the average number of mutations in ORF1ab, S, ORF3a, M and N of genomes, having nucleotide shift at G614 (n=459), were significantly higher (p≤0.001) than those having mutation at D614 (n=215). Previously reported frequent mutations such as P4715L, D614G, R203K, G204R and I300F were also prevalent in Bangladeshi isolates. Additionally, 87 unique amino acid changes were revealed and were categorized as originating from different cities of Bangladesh. The analyses would increase our understanding of variations in virus genomes circulating in Bangladesh and elsewhere and help develop novel therapeutic targets against SARS-CoV-2.

## 1. Introduction

Severe Acute Respiratory Syndrome-CoronaVirus-2 (SARS-CoV-2) has become an etiological agent of the disease called CoronaVirus Disease-2019 (COVID-19). As of 21 August 2020, globally there have been 22,492,312 confirmed cases of COVID-19, including 788,503 deaths, reported to WHO. To explore the viral pathogenesis, modern genomics tools are highly crucial and has been employed by researchers around the world. Hundreds of virus whole genomes are now submitted in publicly accessible databases from different parts of the globe everyday. It is hightime to analyze the variations among those sequences which will help future strategic efforts for its preventive measures such as vaccine design and therapeutics. SARS-CoV-2 consists of positive-sense single-stranded RNA with a genome size ranging from ∼27 to 34 Kb. It contains a variable number (6-11) of open-reading frames (ORFs). The first ORF is almost two-third of the whole genome and encodes four structural, 16 non-structural and eight accessory proteins [1,2]. According to a recent study on 48,635 of SARS CoV-2 genome sequences, 353,341 mutation events have been observed globally in comparison to the reference genome of Wuhan. Among these sequences, India, Congo, Bangladesh and Kazakhstan have significantly high numbers of mutations per sample compared to the global average [3]. Out of these mutations, D614G mutation (causing aspartate to glycine in S protein position 614) is reported to be the most prevalent mutations reported from Europe, Oceania, South America, Africa [4]. Zhang et al. (2020) reported that the level of angiotensin-converting enzyme 2 (ACE2) expression was distinctly higher by the retroviruses pseudotyped with G614 compared to that of D614 [4,5]. The functional properties of the (D614) and (G614) were compared in this study and G614 was found to be more stable than D614 with more transmission efficiency, supporting the previous epidemiological data [5]. Another reported mutation of ORF1ab is P4715L linked with D614G, that has been reported to have a strong relationship with higher fatality rates in 28 countries and 17 states of the United States [3,6]. Three other mutations namely C14408T (ORF1b), C241T (5’ UTR), C3037T (ORF1a) is reported to be common and co-occurring in the same genome while G11083T has been found mostly in Asian countries [6]. Hassan et al. (2020) investigated the accumulation of ORF3a mutations of SARS-CoV-2 from India where they revealed four types of mutations (Q>H, D>Y and S>L) near TRAF, ion channel and caveolin binding domain, respectively [7]. Notable that all these mutations might have implications in maintaining the virulence of the virus and NLRP3 inflammasome activation.

In Bangladesh, as of 21 August, 2020, nearly 287,959 people are infected and 3822 people have died due to COVID-19 (https://iedcr.gov.bd/). Among them, 329 of SARS-CoV-2 genome sequences were deposited in GISAID database (https://www.gisaid.org). Analysis of these sequences is sparsely reported in the literature. For example, Islam et al. (2020) analyzed 60 SARS-CoV-2 genome sequences from Bangladesh and compared them with only six Southeast Asian countries [8]. On the other hand, Parvez et al. (2020) analyzed only 14 isolates and found 42 mutations [9]. Hasan et al. (2020) identified only nine variants where unique mutations (UM) were found prevalent mostly in ORF1ab [10]. However, Shishir et al. (2020) analyzed 64 sequences and identified the presence of 180 mutations in the coding regions of the viruses and mutations at nsp2 was the most prevalent [11]. Due to the small numbers of genome sequence analyzed, most of these findings were not conclusive and representative. Further comprehensive analyses is therefore necessary to better understand the circulating virus in the country.

In this study, we first compared 207 genome sequences isolated from Bangladesh with 467 global sequences mainly from Asia, Africa, Europe, Australia and North America using time-resolved phylogenetic analysis and investigated the origin of imported COVID-19 cases to Bangladesh. Then, we studied the variants present in different isolates of Bangladesh to investigate the pattern of mutations, identify UMs, and discuss the pseudo-effect of these mutations on the structure and function of encoded proteins, with their role in pathogenicity. Most interestingly, we found 87 UMs with a total count of 113 in Bangladeshi isolates which will increase our understanding of distribution of SARS-CoV-2 virus in different regions and associated pathogenicity.

## 2. Methods and materials

### 2.1 Dataset

As of 30 June, 2020, a total of 35,723 whole genomic RNA sequences of SARS-CoV-2 had been submitted to GISAID. From these downloaded sequences, a custom python script was used to retrieve unique sequences. The same script also removed any sequence containing “N” and other ambiguous IUPAC codes [12]. This resulted in a total of 8723 complete genomic sequences. To select representative sequences from curated 8723 sequences and make a comparison against sequences from Bangladesh, priorities were given to those countries that had a higher number of infections in each continent (source: https://www.worldometers.info/coronavirus/). We selected these sequences in such a way that each continent must contain at least one sequence from each GISAID clade. The number of sequences selected from a country was based on the total number of unique sequences retrieved. This resulted in a total of 467 unique representative sequences from 42 countries (see Supporting File Table S1).

### 2.2 Phylogenetic analysis

From Bangladesh, 215 whole-genome RNA sequences of SARS-CoV-2 were uploaded to GISAID as of 15 July, 2020. Only high coverage complete sequences (n=207) were kept for analysis. All these 207 sequences were aligned with the previously selected 467 representative sequences along with that of Wuhan-1 (Accession ID MN908947) as a reference sequence [13]. To ensure comparability, the flanks of all the sequences were truncated to the consensus range from 56 to 29,797 [14], with nucleotide position numbering according to the Wuhan 1 reference sequence, prior to alignment. Multiple Sequence Alignment (MSA) and phylogenetic tree construction were carried out using Molecular Evolutionary Genetic Analysis (MEGA X) software [15]. All selected sequences were aligned using MUSCLE software tool [16]. Later, an NJ (Neighbor-Joining) phylogenetic tree [17] was constructed using the Tamura-Nei method. Tree topology was assessed using a fast bootstrapping function with 1000 replicates. Tree visualization and annotations were performed in the Interactive Tree of Life (iTOL) v5 [18].

### 2.3 Mutation analysis

The Genome Detective Coronavirus Typing Tool Version 1.13 was used for variant analyses of SARS-CoV-2 genome which is specially designed for this virus analysis (https://www.genomedetective.com/app/typingtool/cov/) [19]. For analysis of (UM) among 207 genomic sequences from Bangladesh, we used a CoV server hosted in GISAID server (https://www.gisaid.org/epiflu-applications/covsurver-app/). The server analyzed our dataset against all available genomic sequences of SARS-CoV-2 including the Wuhan reference sequence deposited on GISAID until July 16, 2020.

### 2.4 Statistical analysis

Descriptive and inferential statistics were used to analyze different mutations and their correlation with different categorical variables. For correlation, we used one-way analysis of variance using SPSS Statistics 25 (IBM, Armonk, New York) licensed to King’s College London.

## 3. Results and discussion

### 3.1 Phylogenetic analysis of SARS-CoV-2 circulating in Bangladesh

To understand the SARS-CoV-2 viral transmission in Bangladesh, we performed phylogenetic analysis on the selected 207 viral genomes reported from different districts of Bangladesh along with selected 467 globally submitted sequences as reported from 42 countries and 6 continents (Figure 1). This apparently represents the overall clade distribution of all global sequences along with sequences from Bangladeshi isolates. GR clade was found predominant in Bangladesh as about 84% of the sequences were grouped to this clade followed by GH and G with ∼6 and ∼5%, respectively. Similar clade distribution has been found in isolates submitted from European countries. We also attempted to compare the sequence data among different districts of Bangladesh from where patient samples were collected for sequencing, Looking at the district-wise distribution of clade (Figure 2), it was found that the sequences from three districts, namely Dhaka, Narayanganj and Rangpur primarily belong to GR clade. Conversely, only sequences from Chattogram district were from S and GH clades. On another note, phylogenetic analyses of the clade distribution of isolates from countries like Saudi Arabia, where GH clade was predominant [3] it is highly likely that the introduction of GH and S clades in Bangladesh could be of Middle-Eastern origin. Based on the datsets, we hypothesized that Bangladeshi SARS-CoV-2 isolates belonging to different clades might have critical implications concerning viral transmission rate, virulence, severity and other aspects of disease pathogenesis. In addition, the presence of different clades of SARS-CoV-2 strains in different districts could also have implications in the accuracy of diagnostic tests that are underway.

**Figure 1:**
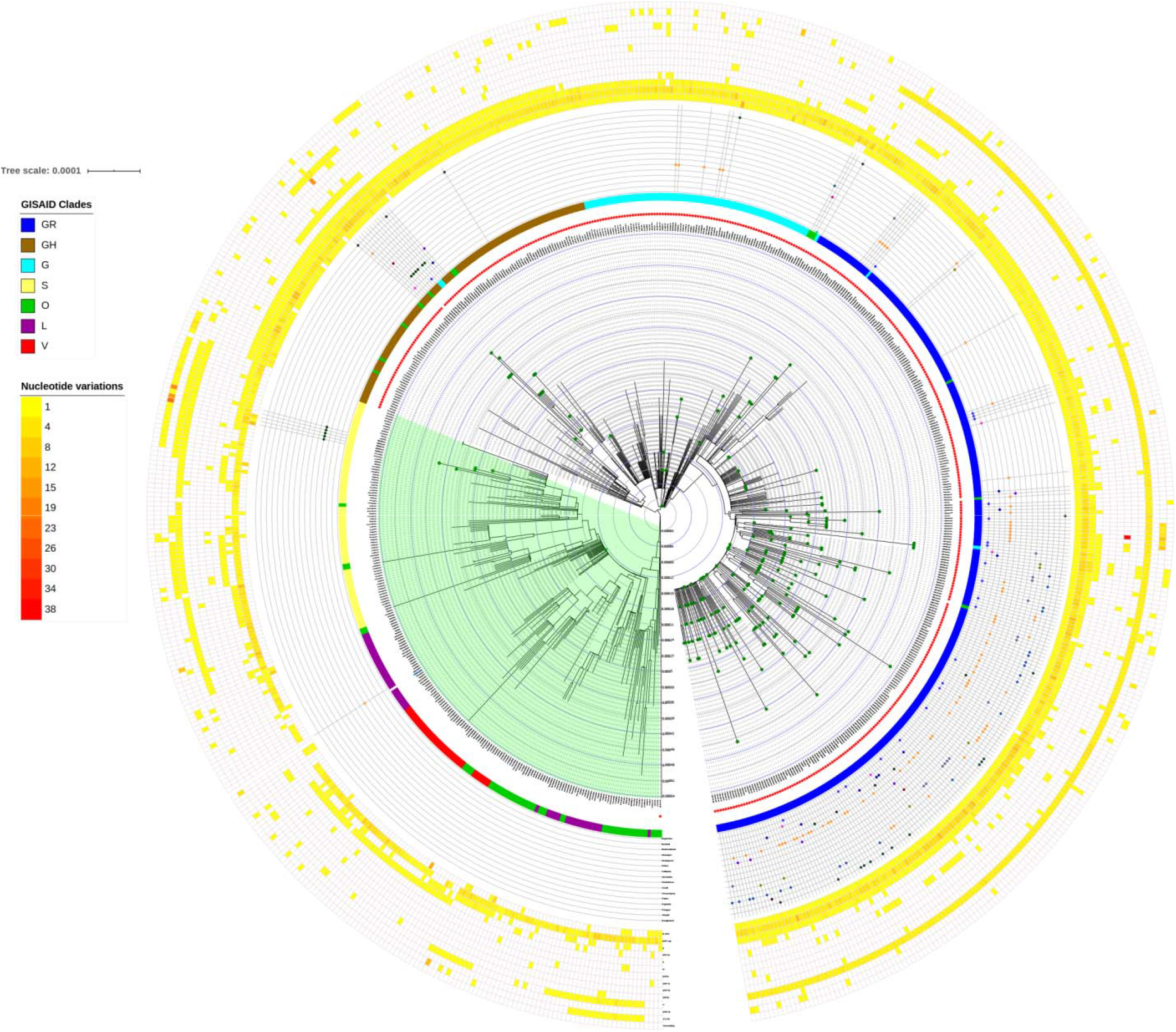
Phylogenetic analysis of 207 Bangladeshi SARS-CoV-2 genomes with 467 representative sequences from 42 different countries worldwide. Sequences from Bangladesh are annotated with ‘green circles’ in the tree and D614G mutations cluster is ‘shaded green’. Outside the main tree there are four different annotation panels sequentially to show the presence of D614G mutations as binary data (G614 as Red Star), followed by GISAID clades as a ‘color strip’, district-wise distribution of Bangladeshi sequences in a ‘shape plot ‘and finally burden of mutation in different ORFs of SARS-CoV-2 in the outermost panel as ‘heatmap’.

**Figure 2:**
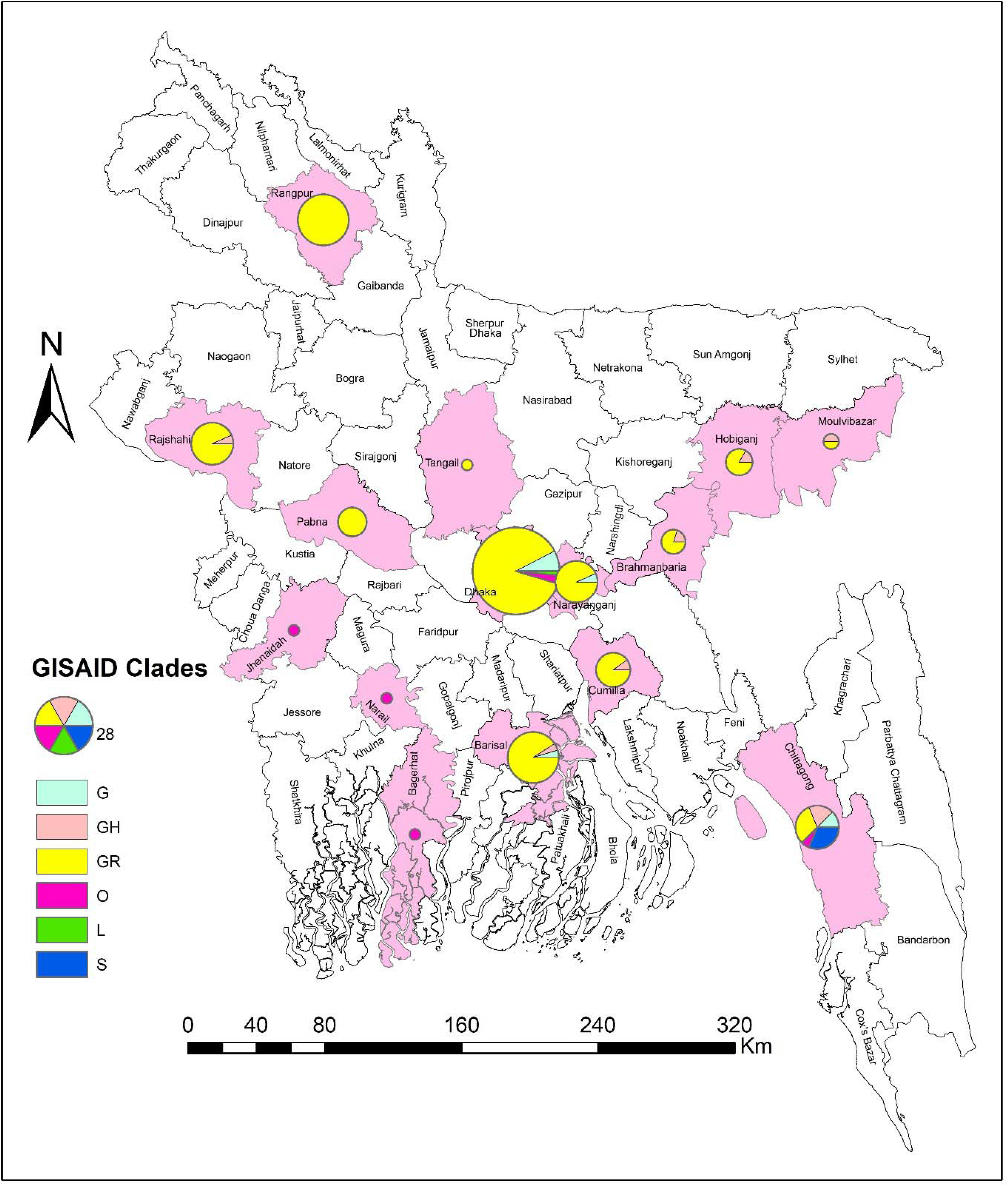
District-wise distribution of GISAID clades among circulating SARS-CoV-2 strains in Bangladesh. Please be noted that district information of 18 sequences out of 207 deposited from Bangladesh is not available from respective metadata. The spatial map was created using layers downloaded from GeoDASH (The Bangladesh Geospatial Data Sharing Platform) website on ArcGIS Desktop (Esri Inc.,109 Redlands, California, United States) licensed to King’s College London.

### 3.2 Mutations analysis in Bangladeshi genomic sequences

#### 3.2.1 Prevalence of global common mutations

Our analyses revealed a total of 2602 mutations observed among 207 Bangladeshi genomic sequences that constitutes a number of 1602 missense, 612 synonymous, 36 insertion/deletions and 352 other mutations (Table 1). We identified mutations like I300F (nsp2), P4715L (nsp12), D614G (S glycoprotein), R203K and G204R (N protein) are the most frequently occurring common mutations found in Bangladesh with a frequency of 138, 199, 198, 177 and 177, respectively (Figure 3). Notable that, no particular mutations occurred at any specific time period rather they have been observed over the whole period of disease incidence. Firstly, 1163A>T (I300F), can be a destabilizing factor for nsp2 and thus modulating the strategy of host cell survival [11,20]. This mutation can lead to a reduction of conformational entropy due to the presence of the side chains that can result in charge neutralization of the phosphorylated serine residues [21]. Secondly, RNA dependent RNA polymerase (nsp12) is significant for replication and transcription of the viral RNA genome. Therefore, P > L at 4715 in nsp12 may have some effects on RNA transcription. This mutation was also observed in most of the USA states (28 out of 31). The same mutation was prevalent in European countries like Spain, France etc. This alteration could affect the pathogenesis triggered by antibody escape variants with the epitope loss [22,23]. Thirdly, Korber et al. (2020) stated that the G614 type might have originated either in Europe or China [24]. They also reported that the original Wuhan D614 form was also predominant in Asian samples. Meanwhile, the G614 form had clearly established and started expanding in countries outside of China. We also noticed 95.6% of genomes from Bangladesh have D614G mutation, which is also dominant in the world. However, the average number of mutations per ORF is varied among D614 and G614 containing genomes that we have studied (n=674) as revealed by Table 2. The average number of mutations in ORF1ab, S, ORF3a, M and N of genomes having mutated G614 (n=459) are significantly higher (*p*≤0.001) than those having wild D614 (n=215) in S glycoprotein (Table 2; Figure 1). Interestingly, the average mutation number is declined in ORF8 of genomes having G614 mutation (*p*<0.001). This correlation indicates that the genomes containing D614G mutation are more prone to bear other mutations which may facilitate the notion that the link of this mutation with the transmission and pathogenesis of SARS-CoV-2 [4,12,25]. Finally, R203K and G204R mutations in N protein were previously reported in Indian, Spanish, Italian, and French samples [21,26]. These mutations are located in the site of the SR-rich region which has been reported to be intrinsically disordered [27]. This region further incorporates a few phosphorylation sites [28], including the GSK3 phosphorylation at Ser202 and a CDK phosphorylation site at Ser206 which are located close to the position of this mutation. The ‘SRGTS’ (202-206) and ‘SPAR’ (206-209) sequence motifs are dependent on GSK3 and CDK phosphorylation motifs, respectively. Other variations (28881G>A and 28882G>A) together convert polar to non-polar amino acid (R203K) and 28883G>C variation converts nonpolar to polar amino acid (G204R).

**Table 1:** Number of common gene variants among 207 SARS-CoV-2 genomes from Bangladesh

| Genome segment | Missense mutation | Synonymous mutation | Insertion Deletion | Others | Total |
| --- | --- | --- | --- | --- | --- |
| ORF1ab | 633 | 458 | 1 | 0 | 1092 |
| S | 250 | 58 | 3 | 0 | 311 |
| ORF3a | 53 | 29 | 5 | 1 | 88 |
| E | 2 | 1 | 0 | 0 | 3 |
| M | 13 | 30 | 0 | 0 | 43 |
| ORF6 | 3 | 1 | 0 | 0 | 4 |
| ORF7a | 35 | 4 | 9 | 0 | 48 |
| ORF7b | 1 | 1 | 0 | 0 | 2 |
| ORF8 | 30 | 7 | 2 | 0 | 39 |
| ORF10 | 3 | 5 | 0 | 0 | 8 |
| N | 579 | 17 | 3 | 0 | 599 |
| Intergenic | 0 | 0 | 2 | 8 | 10 |
| 5' UTR | 0 | 1 | 4 | 230 | 235 |
| 3' UTR | 0 | 0 | 7 | 113 | 120 |
| **Total** | **1602** | **612** | **36** | **352** | **2602** |

**Table 2:** Correlation of the average number of mutations per genome among different genomic segments with D614G mutation.

| Genome segment | Average number of mutations per genome |  | Change (%) | *p* value |
| --- | --- | --- | --- | --- |
|  | D614 (wild; n=215) | G614 (n=459) |  |  |
| 5' UTR | 0.37 | 1.12 | 67 | 0.000 |
| ORF1ab | 3.23 | 4.36 | 26 | 0.000 |
| S | 0.52 | 1.39 | 63 | 0.000 |
| ORF3a | 0.40 | 0.44 | 7.0 | 0.530 |
| E | 0.00 | 0.01 | 57 | 0.422 |
| M | 0.07 | 0.17 | 58 | 0.001 |
| ORF6 | 0.01 | 0.03 | 64 | 0.153 |
| ORF7a | 0.02 | 0.12 | 84% | 0.417 |
| ORF7b | 0.01 | 0.01 | -42 | 0.697 |
| ORF8 | 0.53 | 0.06 | -809 | 0.000 |
| N | 0.60 | 1.96 | 70 | 0.000 |
| ORF10 | 0.01 | 0.02 | 36 | 0.491 |
| 3' UTR | 0.69 | 0.27 | -155 | 0.002 |
| Total | 6.48 | 9.98 | 35 |  |

**Figure 3:**
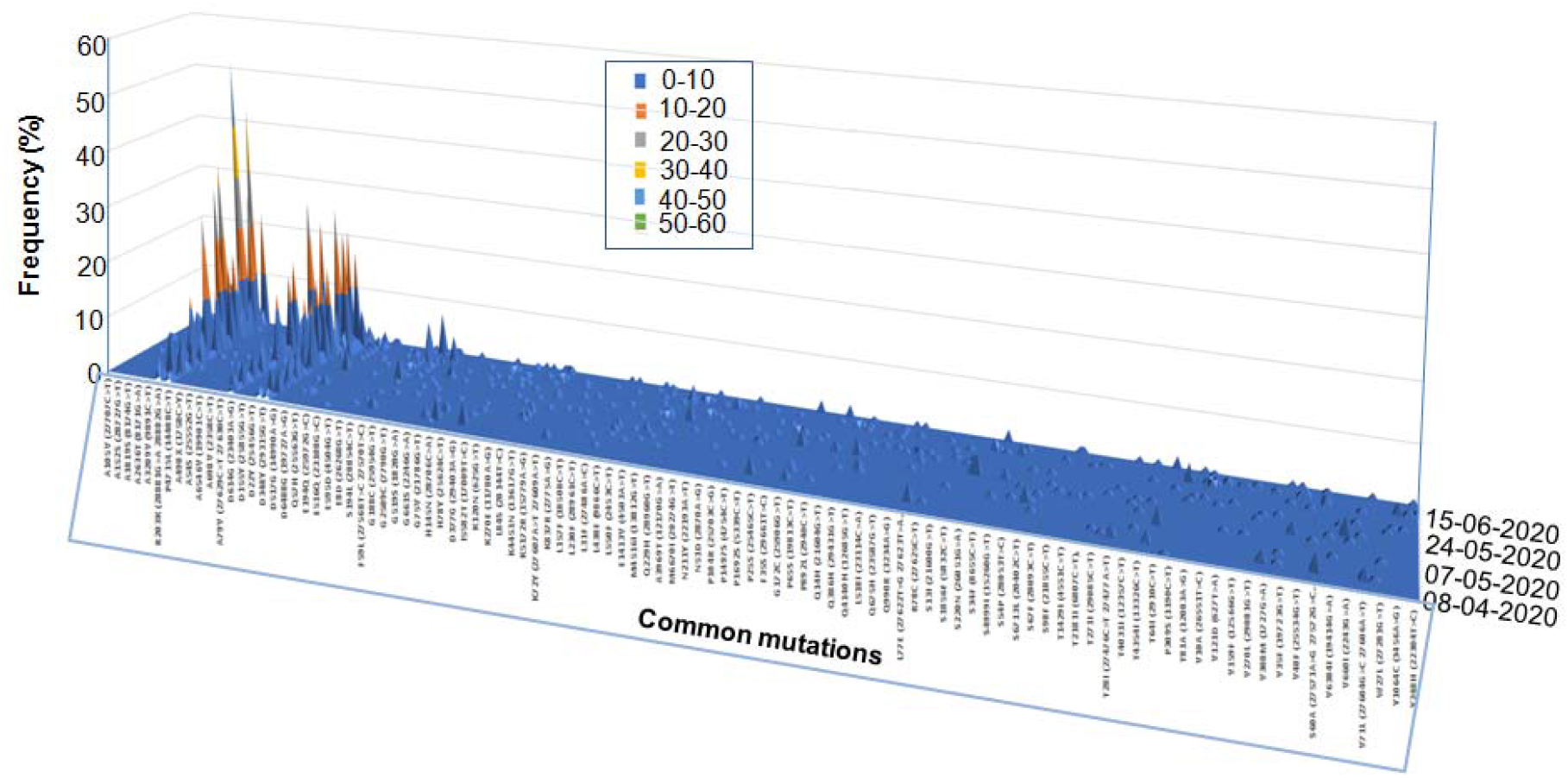
Frequency of common mutations observed in Bangladesh over time.

#### 3.2.2 Unique mutations (UM) analysis in Bangladeshi isolates

We observed 87 unique shifts from the different proteins of SARS-CoV-2 genomes found in Bangladesh (Table S2). Details of their pseudo-effects on viral replication, assembly, transmissivity and pathogenicity are corroborated in Table S2. Surprisingly, most of these UM were localized except a few exceptions. For example, the UMs E8D (E), S180T (N), L139J and S220N (NS3), S54P (NS8), V56A (nsp1), A550V and S607I (nsp12), Q470R and Y198H (nsp13), V437F (nsp14), D409B and N377D (nsp2), T1363I and Y246C (nsp3), D85E (nsp4), N133B and R188S (nsp5) and S939Y (S) were only found in Dhaka, which is the capital city of Bangladesh. The second most important city in Bangladesh is Chattogram where many UMs were found, such as G254stop (NS3), N39Y (NS6), L4I, L7G and V5T (NS8), T101I, (nsp10), V459I (nsp14), A602S, N1337S and I1672S (nsp3). Similarly, Rangpur has UMs of F20L (E), Q83R (N), W69R (NS3), G42V (NS7), Q224K (nsp12), I258T (nsp13), R883G, S1038F and Y272H (nsp3). Barishal also shows UMs of E194Q and G188C (NS3), Q62E (NS7a), A529S, D517G and E254D (nsp12), L22I (nsp6) and T81A (nsp7). A few UMs have been observed in some other districts like Chandpur, Pabna, Rajshahi, Tangail, Pabna and Brahmanbaria. These changes in amino acids might have occurred due to rapid mutation and/or recombination with existing other CoV in the human body [7]. Another possibility of these mutations is the zoonotic origin of the SARS-CoV-2 in Bangladesh. Interestingly, some districts shared UMs, such as Q38E (Rajshahi, Pabna and Barishal), S220N (Dhaka and Rajshahi), V255del (Narayanganj and Barishal) in NS3; P38R (Brahmanbaria, Chandpur and Pabna) in NS8; V469A (Chattogram and Barishal), V594F (Brahmanbaria, Chandpur and Pabna) in nsp2; L373M (Brahmanbaria, Rajshahi and Moulvibazar); N51D (Dhaka, Chattagram and Barishal) in nsp3; I106S (Chandpur and Hobigonj) in nsp5; K270E (Dhaka and Barishal), V120L (Pabna and Hobigonj) in nsp6; and F140del (Rangpur and Dhaka), L518I (Rajshahi, Brahmanbaria and Moulvibazar), N211Y (Dhaka and Chandpur) in S protein. Circulation of a high number of these UMs in different cities, indicates the possible emergence of community transmission in Bangladeshi population [29].

Among 87 amino acids mutations, 66 mutations were located in nsps-1 (4), 2 (9), 3 (22), 4 (3), 5 (5), 6(5), 7(1), 10 (1), nsp-12 (7), 13 (3), 14(2) and 15 (4) at ORF1ab. These 66 UMs constitute around 60% of the total UMs found which imply most of the change of SARS-CoV-2’s virulence property in Bangladesh patients. ORF1ab encodes a multifunctional polyprotein which is involved in the transcription and replication of CoV RNAs. We observed 4 UMs in nsp1, but the location of these amino acids was not in the KH domain (K164 and H165 of nsp1), which binds with ribosome 40s subunit [30]. However, nsp1 acts as a primary virulence factor in SARS-CoV-2 infection, and mutation in this protein could affect the structure and functional properties, thereby altering its virulence properties. Nine UMs were seen in nsp2, but their effects on host cells were merely reported in the literature. Since nsp2 interacts with the host proteins and disrupts the host cell survival signaling pathway [31], any mutation in nsp2 may play a crucial role in SARS-CoV infections. In a recent study, it has been found that, compared to Bat SARS-CoV, SARS-CoV-2 has a stabilizing mutation at amino acid position 501, T501Q, which alters the viral pathogenicity and makes the virus more contagious [32]. We did not observe this mutation in our study. The mutations of nsp3 are responsible for affecting the virus assembly and hence their replication. It is due to the disruption of replicase polyprotein processing into nsps. These nsps assemble with cellular membranes and facilitate virus replication. The UMs found in nsp3 might have some probable effects. Firstly, the UMs (A889V, R883G, S1038F and V843F) were found in the main domain of nsp3 that is important for processing endopeptidases from coronaviruses. Secondly, many UM (e.g. A1803V, A1819S, G1691C, I1672S, A602S, D782B, K462R, L373M, N1337S, N51D, R883G, T1363Is, Y246C and Y272H etc) are found in topological (cytoplasmic) domain rather than transmembrane domain which could interfere less on cytoplasmic double-membrane vesicle formation, necessary for viral replication. Thirdly, we observed three UMs (L373M, Y246C and Y272H) in ADP-ribose-1′-phosphatase (ADRP) or (Macro) domain. It has been shown that mutations of the ADRP domains does not diminish virus replication in mice, but reduces the production of the cytokine IL-6, which is an important pro-inflammatory molecule [33]. However, we did not observe any mutation in the active sites and zinc finger motif, attributing normal catalytic activity of nsp3. The general opinion is that SARS-CoV PL-PRO domain is important for the development of antiviral drugs and of the actual role of this enzyme in the biogenesis of the COVID-19 replicase complex is yet not explored. It is proposed that the proteins nsp3, nsp4 and nsp6, through their transmembrane domains, are involved in the replicative and transcription complex [34]. In our study, we observed only 3 UMs in nsp4 and nsp6, respectively. Meanwhile, nsp5 encodes 3C-like proteinase which cleaves the C-terminus of replicase polyprotein at 11 sites. The five UMs that we found in nsp5 did not fall in its active sites (3304 and 3408 in ORF1ab) requiring further investigation with large number of sequence datasets.

## 4. Conclusion

The present global outbreak of COVID-19, caused by SARS-CoV-2, has already taken ∼0.8 million lives. To combat this deadly disease, we need a greater understanding of the pathobiology of the virus. Hence, it is essential to minimize the translational gap between viral genomic information and its clinical consequences for developing effective therapeutic strategies. In this study, we have attempted to explore genomic variations of Bangladeshi SARS-CoV-2 viral isolates while comparing with a large cohort of global isolates. Our analyses will facilitate the understanding of the origin, mutation patterns and their possible effect on viral pathogenicity. This study tries to address the importance of the variations in the viral genomes and their necessity for therapeutic interventions. The unique insights from this study will undoubtedly be supportive for a better understanding of SARS-CoV-2 molecular mechanism and to draw an end to the current life-threatening pandemic.

## Notes

### Competing Interest Statement

The authors have declared no competing interest.

