## Supporting Files Table S1 and S2 for "Global and Local Mutations in Bangladeshi SARS-CoV-2 Genomes"

**Table S1**: List of countries and the number of sequences included in this study

| Country | GISAID Clades | | | | | | | Total |
| --- | --- | --- | --- | --- | --- | --- | --- | --- |
|  | G | GH | GR | L | O | S | V |  |
| Argentina | 4 | 4 | 6 | 0 | 0 | 0 | 0 | 14 |
| Australia | 6 | 7 | 7 | 3 | 2 | 7 | 2 | 34 |
| Bangladesh | 10 | 12 | 173 | 1 | 6 | 5 | 0 | 207 |
| Benin | 0 | 0 | 0 | 0 | 0 | 2 | 0 | 2 |
| Brazil | 4 | 2 | 6 | 1 | 4 | 2 | 2 | 21 |
| Canada | 2 | 5 | 7 | 2 | 3 | 7 | 0 | 26 |
| Chile | 5 | 7 | 6 | 2 | 0 | 4 | 1 | 25 |
| China | 2 | 3 | 3 | 2 | 2 | 3 | 2 | 17 |
| Colombia | 5 | 5 | 6 | 0 | 0 | 2 | 0 | 18 |
| Democratic Republic of the Congo | 5 | 2 | 3 | 0 | 0 | 2 | 1 | 13 |
| Egypt | 0 | 5 | 0 | 0 | 0 | 1 | 0 | 6 |
| France | 2 | 3 | 2 | 1 | 5 | 1 | 0 | 14 |
| Gambia | 0 | 0 | 1 | 0 | 0 | 0 | 0 | 1 |
| Germany | 0 | 1 | 2 | 2 | 1 | 1 | 1 | 8 |
| Hong Kong | 1 | 0 | 0 | 1 | 2 | 1 | 0 | 5 |
| India | 2 | 2 | 2 | 2 | 3 | 2 | 0 | 13 |
| Indonesia | 0 | 0 | 0 | 3 | 1 | 0 | 0 | 4 |
| Iran | 0 | 0 | 0 | 1 | 1 | 0 | 0 | 2 |
| Italy | 3 | 0 | 3 | 0 | 0 | 0 | 2 | 8 |
| Japan | 0 | 2 | 2 | 2 | 0 | 0 | 0 | 6 |
| Malaysia | 0 | 0 | 0 | 2 | 2 | 1 | 0 | 5 |
| Mexico | 2 | 1 | 3 | 0 | 0 | 7 | 0 | 13 |
| Morocco | 5 | 0 | 0 | 0 | 0 | 0 | 0 | 5 |
| Nigeria | 0 | 2 | 0 | 0 | 0 | 4 | 1 | 7 |
| Oman | 1 | 1 | 1 | 1 | 1 | 0 | 1 | 6 |
| Pakistan | 0 | 2 | 0 | 0 | 1 | 1 | 0 | 4 |
| Russia | 4 | 7 | 7 | 1 | 0 | 1 | 1 | 21 |
| Saudi Arabia | 1 | 5 | 0 | 0 | 2 | 3 | 0 | 11 |
| Senegal | 2 | 0 | 2 | 0 | 0 | 0 | 0 | 4 |
| Singapore | 1 | 2 | 2 | 3 | 2 | 2 | 2 | 14 |
| South Africa | 1 | 0 | 0 | 0 | 0 | 0 | 0 | 1 |
| South Korea | 0 | 0 | 0 | 0 | 1 | 2 | 4 | 7 |
| Spain | 2 | 3 | 3 | 2 | 5 | 5 | 4 | 24 |
| Sri Lanka | 1 | 0 | 1 | 0 | 2 | 0 | 0 | 4 |
| Sweden | 2 | 2 | 2 | 6 | 7 | 2 | 1 | 22 |
| Taiwan | 1 | 1 | 1 | 0 | 1 | 1 | 1 | 6 |
| Thailand | 0 | 1 | 1 | 0 | 0 | 1 | 0 | 3 |
| Turkey | 0 | 0 | 1 | 0 | 3 | 0 | 0 | 4 |
| United Arab Emirates | 2 | 2 | 2 | 2 | 2 | 1 | 0 | 11 |
| United Kingdom | 3 | 2 | 3 | 4 | 0 | 0 | 3 | 15 |
| USA | 6 | 6 | 6 | 5 | 6 | 6 | 5 | 40 |
| Vietnam | 0 | 1 | 1 | 0 | 0 | 0 | 1 | 3 |
| **Grand Total** | **85** | **98** | **265** | **49** | **65** | **77** | **35** | **674** |

**Table S2:** List of unique mutations and their possible role in SARS-CoV-2 pathogenicity

| Protein | Mutation | Possible role or impact | District |
| --- | --- | --- | --- |
| ***ORF1ab*** |  |  | |
| NSP1 | V56A (432T>C) | Hydrophobic Val to simple, non-polar Ala | Dhaka |
|  | M85del (518_520delATG) | Deletion of Met | Unspecified |
|  | V121D (627T>A) | Hydrophobic Val to acidic Asp | Unspecified |
|  | L122I (629C>A) | The same type of change just increased in size | Unspecified |
| NSP2 | L71S (1017T>C) | Hydrophobic side chain containing Leu to polar, non-charged Ser | Rajshahi |
|  | V308M (1727G>A) | Hydrophobic Val to sulfur containing Met | Chandpur |
|  | N377D (1934A>G) | Polar Asn to acidic Asp | Dhaka |
|  | D409B (Ambiguous) |  | Dhaka |
|  | V469A (2211T>C) | Hydrophobic Val to another hydrophobic, non-polar Ala | Chattogram, Barishal |
|  | V594F (2585G>T) | Hydrophobic Val to aromatic Phe | Brahmanbaria, Chandpur, Pabna |
| NSP3 | N51D (2870A>G) | Polar aliphatic Asn to acidic Asp | Dhaka, Chattogram, Barishal |
|  | Y246C (3456A>G | Aromatic Tyr to sulfur containing acidic Cys | Dhaka, Rajshahi |
|  | Y272H (3533T>C) | Aromatic Tyr to polar, basic His | Rangpur |
|  | L373M (3836T>A | Hydrophobic Leu to sulfur containing, hydrophobic Met | Rajshahi, Brahmanbaria, Moulvibazar |
|  | K462R (4104A>G) | Positively charged ε-amino group containing Lys to positively charged Arg | Rajshahi |
|  | A602S (4523G>T) | Hydrophobic, non-polar Ala to hydrophilic Ser | Chattogram |
|  | D782B (Ambiguous) | Ambiguous | Unspecified |
|  | V843F (5246G>T) | Non-polar/hydrophobic Val to aromatic Phe | Unspecified |
|  | R883G (5366A>G) | Positively charged Arg to non-polar Gly | Rangpur |
|  | A889V (5385C>T) | Same change, just larger in size | Unspecified |
|  | S1038F (5832C>T) | Hydrophilic Ser to aromatic Phe | Rangpur |
|  | N1337S (6729A>G) | Polar Asn to hydrophilic Ser | Chattogram |
|  | T1363I (6807C>T | Hydrogen bonding Thr to non-polar/hydrophobic Ile | Dhaka |
|  | I1672S (7734T>G) | Non-polar/hydrophobic Ile to hydrophilic Ser | Chattogram |
|  | G1691C (7790G>T) | Simple Gly to sulfur-containing, acidic Cys | Unspecified |
|  | A1803V (8127C>T) | Same change, just larger in size | Rangpur |
|  | A1819S (8174G>T) | Simple, hydrophobic Ala to polar, non-charged Ser | Pabna |
| NSP4 | A69V (8760C>T) | Same change, just larger in size | Pabna |
|  | D85E (8809C>A) | Both are the same type of amino acid | Dhaka |
|  | E425G (9828A>G) | Acidic Glu to hydrophobic Gly | Hobigonj |
| NSP5 | P96S (10340C>T) | Bulky, hydrophobic phenylalanine to polar, non-charged Ser | Chandpur |
|  | I106S (10371T>G) | Hydrophobic Ile to polar Ser | Chandpur, Hobigonj |
|  | N133B (Ambiguous) |  | Dhaka |
|  | R188S (10618G>T) | Positively charged Arg to polar Ser | Dhaka |
|  | L22I (11036C>A) | Both are the same type of amino acids | Barishal |
| NSP6 | V120L (11330G>T) | Valine to hydrophobic side chain containing Leu | Pabna, Hobigonj |
|  | K270E (11780A>G) | Positively charged ε-amino group containing Lys to negatively charged Glutamic acid | Dhaka, Barishal |
| NSP7 | T81A  (12083A>G) | Hydroxyl group containing Thr to hydrophobic Ala | Barishal |
| NSP10 | T101I (13326C>T) | Hydrogen bonding Thr to hydrophobic Ile | Chattogram |
| NSP12 | Q224K (14110C>A) | Neutral Gln to positively charged ε-amino group containing Lys | Rangpur |
|  | E254D (14202G>T) | Acidic Glu to another acidic Asp | Barishal |
|  | D517G (14990A>G) | Acidic Asp to hydrophobic Gly | Barishal |
|  | A529S (15025G>T) | Hydrophobic Ala to polar Ser | Barishal |
|  | A550V (15089C>T) | Same change, just larger in size | Dhaka |
|  | S607I (15260G>T) | Polar Ser to hydrophobic Ile | Dhaka, Dhaka |
| NSP13 | Y198H (16828T>C) | Aromatic Tyr to polar, basic His | Dhaka |
|  | I258T (17009T>C) | Hydrophobic side chain containing amino acid Ile to hydroxyl group containing Thr | Rangpur |
|  | Q470R (17645A>G) | Neutral Gln to positively charged Arg | Dhaka |
| NSP14 | V437F (19348G>T) | Aliphatic, hydrophobic Val to non-polar, aromatic Phe | Dhaka |
|  | V459I (19414G>A) | Aliphatic, hydrophobic Val to hydrophobic Ile (larger in size) | Chattogram |
| NSP15 | D36G (19727A>G) | Acidic Asp to hydrophobic Gly | Tangail |
|  | E68D (19824G>T) | Acidic Glu to another acidic Asp | Chandpur |
|  | A94V (19901C>T) | Same change, just larger in size | Unspecified |
| ***ORFS*** |  |  | |
| S Glycoprotein | F140del (21980_21982delTTT) | Deletion of Phe | Rangpur, Dhaka |
|  | Y145del (21992_21994delTAT) | Deletion of Tyr | Unspecified |
|  | N211Y (22193A>T) | Polar aliphatic Asn to Aromatic, neutral and hydrophobic Tyr | Dhaka, Chandpur |
|  | Y248H (22304T>C) | Aromatic Tyr to polar, basic His | Barishal |
|  | E516Q (23108G>C) | Negatively charged Glu to neutral Gln | Narail |
|  | L518I (23114C>A) | Both are hydrophobic side chain containing amino acid. Differ in size | Rajshahi, Brahmanbaria, Moulvibazar |
|  | E654Q (23522G>C) | Negatively charged Glu to neutral Gln | Brahmanbaria |
|  | Y660F (23541A>T) | Aromatic Tyr to Aromatic Phe | Chandpur |
|  | S939Y (24378C>A) | Polar, non-charged Ser to Aromatic Tyr | Dhaka |
| ***ORF3*** |  |  | |
| NS3 | Q38E (25504C>G) | Neutral Gln to negatively charged Glu | Unspecified, Unspecified, Rajshahi, Rajshahi, Pabna, Barishal |
|  | W69R (25597T>C) | Aromatic Trp to positively charged Arg | Rangpur |
|  | P104R (25703C>G) | Bulky, hydrophobic phenylalanine to positively charged Arg | Unspecified |
|  | L139J (Ambiguous) |  | Dhaka |
|  | G188C (25954G>T) | Hydrophobic Gly to sulfur-containing, acidic Cys | Barishal |
|  | E194Q (25972G>C) | Negatively charged Glu to neutral Gln | Barishal |
|  | S220N (26051G>A) | Polar, non-charged Ser to polar Asn | Dhaka, Rajshahi |
|  | G254stop (26152G>T) | Hydrophobic Gly to stop codon | Chattogram |
|  | V255del (26159_26161delTTA) | Hydrophobic Val is deleted | Narayanganj, Barishal, Barishal, Barishal |
| ***ORFE*** |  |  | |
| E | E8D (26268G>T) | Glutamic acid converted to aspartic acid. Aspartic has one less methylene group than glutamic acid | Dhaka |
|  | F20L (26302T>C) | Phenylalanine convert to leucine which doesn’t have benzene ring | Rangpur |
| ***ORF6*** |  |  | |
| NS6 |  |  | |
|  | N39Y (27316A>T) | Asparagine which contain polar R group convert to Tyrosine which contains aromatic R group | Chattogram |
| ***ORF7a*** |  |  | |
| NS7a | G42V (27518G>T 27519C>T) | Glycine convert to hydrophobic valine | Rangpur |
|  | Q62E (27577C>G) | Glutamine converts to negatively charged glutamic acid | Barishal |
| ***ORF8*** |  |  | |
| NS8 | L4I (27903C>A) | The same type of amino acid, just change in size | Chattogram |
|  | V5T (27906G>A 27907T>C 27908T>G) | Hydrophobic valine convert to Threonine which has polar side chain. | Chattogram |
|  | L7G (27912T>G 27913T>G 27914A>T) | Leucine contains hydrophobic side chains where glycine doesn't contain any side chain. | Chattogram |
|  | P38R (28006C>G) | Phenylalanine convert to Arginine which contains the positively charged group. | Brahmanbaria, Chandpur, Pabna |
|  | S54P (28053T>C) | Ser converted to bulky, hydrophobic Phe | Dhaka |
| ***ORFN*** |  |  | |
| N | D3Y (28280G>T) | Acidic Asp to aromatic Tyr | Pabna |
|  | Q83R (28521A>G) | Glutamine converts to positively charged Arg | Rangpur |
|  | H145N (28706C>A | Hydrophilic His to polar aliphatic Asn | Unspecified |
|  | S180T (28812G>C) | Polar, non-charged Ser to hydroxyl group containing Thr | Dhaka |
| ***ORF10*** |  |  | |
| NS10 | T101I (13326C>T) | Hydroxyl group containing Thr to hydrophobic Ile | Chattogram |
